## Supplementary material for "A structural mechanism for phosphorylation-dependent inactivation of the AP2 complex": Key Resources Table

| Reagent type (species) or resource | Designation | Source or reference | Identifiers | Additional information |
|---|---|---|---|---|
| gene (*Caenorhabditis elegans* ) | *ncap-1* | NA | CELE_Y110A2AR.3 | |
| gene (*C. elegans* ) | *fcho-1* | NA | CELE_F56D12.6 | |
| gene (*C. elegans* ) | *apa-2* | NA | CELE_T20B5.1 | |
| gene (*C. elegans* ) | *apb-1* | NA | CELE_Y71H2B.10 | |
| gene (*C. elegans* ) | *apm-2* | NA | CELE_R160.1 | |
| strain, strain background (*C. elegans* , hermaphrodite) | N2 | PMID: 4366476 | RRID:WB-STRAIN:N2_(ancestral) | Wild type |
| strain, strain background (*C. elegans* , hermaphrodite) | GUN55 | DOI: 10.7554/eLife.32242.001 | | *mewSi81[Pdpy-30:apb-1(trunk):GFP:unc-54(3'UTR)] I; fcho-1(ox477::unc-119(+)) mewSi2[Pdpy-30:RFP:NCAP-1:unc-54(3'UTR)] ncap-1(mew39[1.4 kb deletion]) II; apm-2(ox562[E306K]+mew46[T160A])X* |
| strain, strain background (*C. elegans* , hermaphrodite) | GUN61 | DOI: 10.7554/eLife.32242.001 | | *mewSi1[Pdpy-30:apb-1(trunk):GFP:unc-54(3'UTR)] I; fcho-1(ox477::unc-119(+)) mewSi2[Pdpy-30:RFP:NCAP-1:unc-54(3'UTR)] ncap-1(mew39[1.4 kb deletion]) II* |
| strain, strain background (*C. elegans* , hermaphrodite) | GUN62 | DOI: 10.7554/eLife.32242.001 | | *mewSi81[Pdpy-30:apb-1(trunk):GFP:unc-54(3'UTR)] I; fcho-1(ox477::unc-119(+)) mewSi2[Pdpy-30:RFP:NCAP-1:unc-54(3'UTR)] ncap-1(mew39[1.4 kb deletion]) II; apm-2(ox562[E306K])X* |
| strain, strain background (*C. elegans* , hermaphrodite) | GUN87 | This paper | | *ncap-1(mew54[ncap-1:mScarlet:3xFLAG]) II* |
| strain, strain background (*C. elegans* , hermaphrodite) | GUN88 | This paper | | *ncap-1(mew54[ncap-1:mScarlet:3xFLAG]) fcho-1(ox477::unc-119(+)) II* |
| strain, strain background (*C. elegans* , hermaphrodite) | GUN89 | This paper | | *fcho-1(ox477::unc-119(+)) oxSi883[Phsp-16.41::TEVprotease unc-119(+)] ncap-1(mew39[1.4 kb deletion]) II; mewSi12[Pdpy-30:RFP:NCAP-1(1-163)] IV; apm-2(ox546[W64X]) oxSi877[Papm-2::3xFLAG:APM-2:tev-site unc-119(+)] X* |
| strain, strain background (*C. elegans* , hermaphrodite) | GUN106 | DOI: 10.7554/eLife.32242.001 | | *fcho-1(ox477::unc-119(+)) oxSi883[Phsp-16.41::TEVprotease unc-119(+)] ncap-1(mew39[1.4 kb deletion]) II; mewSi3[Pdpy-30:RFP:NCAP-1:unc-54(3'UTR)] IV; apm-2(ox546[W64X]) oxSi877[Papm-2::3xFLAG:APM-2:tev-site unc-119(+)] X* |
| strain, strain background (*C. elegans* , hermaphrodite) | GUN123 | DOI: 10.7554/eLife.32242.001 | | *mewSi81[Pdpy-30:apb-1(trunk):GFP:unc-54(3'UTR)] I; fcho-1(ox477::unc-119(+)) mewSi32[Pdpy-30:RFP:NCAP-1(S84N):unc-54(3'UTR)] ncap-1(mew39[1.4 kb deletion]) II; apm-2(ox562[E306K])X* |
| strain, strain background (*C. elegans* , hermaphrodite) | GUN135 | DOI: 10.7554/eLife.32242.001 | | *fcho-1(ox477::unc-119(+)) oxSi883[Phsp-16.41::TEVprotease unc-119(+)] ncap-1(mew39[1.4 kb deletion]) II; mewSi35[Pdpy-30:RFP:NCAP-1(S84N)] IV; apm-2(ox546[W64X]) oxSi877[Papm-2::3xFLAG:APM-2:tev-site unc-119(+)] X* |
| strain, strain background (*C. elegans* , hermaphrodite) | GUN195 | This paper | | *fcho-1(ox477::unc-119(+)) oxSi883[Phsp-16.41::TEVprotease unc-119(+)] ncap-1(mew39[1.4 kb deletion]) II; mewSi37[Pdpy-30:RFP:NCAP-1(1-129)] IV; apm-2(ox546[W64X]) oxSi877[Papm-2::3xFLAG:APM-2:tev-site unc-119(+)] X* |
| strain, strain background (*C. elegans* , hermaphrodite) | GUN196 | This paper | | *fcho-1(ox477::unc-119(+)) oxSi883[Phsp-16.41::TEVprotease unc-119(+)] ncap-1(mew39[1.4 kb deletion]) II; mewSi38[Pdpy-30:RFP:NCAP-1(130-236)] IV; apm-2(ox546[W64X]) oxSi877[Papm-2::3xFLAG:APM-2:tev-site unc-119(+)] X* |
| strain, strain background (*C. elegans* , hermaphrodite) | GUN198 | This paper | | *fcho-1(ox477::unc-119(+)) oxSi883[Phsp-16.41::TEVprotease unc-119(+)] ncap-1(mew39[1.4 kb deletion]) II; mewSi40[Pdpy-30:RFP:unc-54(3'UTR)] IV; apm-2(ox546[W64X]) oxSi877[Papm-2::3xFLAG:APM-2:tev-site unc-119(+)] X* |
| strain, strain background (*C. elegans* , hermaphrodite) | GUN204 | This paper | | *mewSi81[Pdpy-30:apb-1(trunk):GFP:unc-54(3'UTR)] I; fcho-1(ox477::unc-119(+)) mewSi46[Pdpy-30:RFP:unc-54(3'UTR)] ncap-1(mew39[1.4 kb deletion]) II; apm-2(ox562[E306K])X* |
| strain, strain background (*C. elegans* , hermaphrodite) | GUN249 | This paper | | *mewSi81[Pdpy-30:apb-1(trunk):GFP:unc-54(3'UTR)] I; fcho-1(ox477::unc-119(+)) mewSi62[Pdpy-30:RFP:NCAP-1:unc-54(3'UTR)(R109E)] ncap-1(mew39[1.4 kb deletion]) II; apm-2(ox562[E306K])X* |
| strain, strain background (*C. elegans* , hermaphrodite) | GUN252 | This paper | | *fcho-1(ox477::unc-119(+)) oxSi883[Phsp-16.41::TEVprotease unc-119(+)] ncap-1(mew39[1.4 kb deletion]) II; mewSi65[Pdpy-30:RFP:NCAP-1(R109E)] IV; apm-2(ox546[W64X]) oxSi877[Papm-2::3xFLAG:APM-2:tev-site unc-119(+)] X* |
| strain, strain background (*C. elegans* , hermaphrodite) | GUN252 | This paper | | *mewSi1[apb-2(trunk):GFP] I; fcho-1(ox477::unc-119(+)) mewSi66[Pdpy-30:RFP:NCAP-1:unc-54(3'UTR)(K148A,E149A,G150A)] ncap-1(mew39[1.4 kb deletion]) II; apm-2(ox562[E306K])X* |
| strain, strain background (*C. elegans* , hermaphrodite) | GUN256 | This paper | | *mewSi81[Pdpy-30:apb-1(trunk):GFP:unc-54(3'UTR)] I; fcho-1(ox477::unc-119(+)) mewSi2[Pdpy-30:RFP:NCAP-1:unc-54(3'UTR)] ncap-1(mew39[1.4 kb deletion]) II; apa-2(mew126[V469D]) apm-2(ox562[E306K])X* |
| strain, strain background (*C. elegans* , hermaphrodite) | GUN271 | This paper | | *mewSi82[apb-1(trunk, S444P)] I; fcho-1(ox477::unc-119(+)) mewSi2[Pdpy-30:RFP:NCAP-1:unc-54(3'UTR)] ncap-1(mew39[1.4 kb deletion]) II; apb-1(mew138[S444P]) III; apm-2(ox562[E306K])X* |
| strain, strain background (*C. elegans* , hermaphrodite) | GUN276 | This paper | | *fcho-1(ox477::unc-119(+)) oxSi883[Phsp-16.41::TEVprotease unc-119(+)] ncap-1(mew39[1.4 kb deletion]) II; mewSi75[Pdpy-30:RFP:NCAP-1(1-190)] IV; apm-2(ox546[W64X]) oxSi877[Papm-2::3xFLAG:APM-2:tev-site unc-119(+)] X* |

| | | | | |
|---|---|---|---|---|
| strain, strain background (*C. elegans* , hermaphrodite) | GUN288 | This paper | | ncap-1(mew54[ncap-1:mScarlet:3xFLAG]) fcho-1(ox477) II; apb-1(mew150[S444P]) III |
| strain, strain background (*C. elegans* , hermaphrodite) | GUN291 | This paper | | ncap-1(mew54[ncap-1:mScarlet:3xFLAG]) fcho-1(ox477::unc-119(+)) II; apa-2(mew153[V469D]) X |
| strain, strain background (*C. elegans* , hermaphrodite) | GUN294 | This paper | | fcho-1(mew55) ncap-1(mew54[ncap-1:mScarlet:3xFLAG]) II apm-2(mew156[T160A]) X |
| strain, strain background (*C. elegans* , hermaphrodite) | GUN296 | This paper | | fcho-1(ox477::unc-119(+)) oxSi883[Phsp-16.41::TEVprotease unc-119(+)] ncap-1(mew39[1.4 kb deletion]) II; mewSi3[Pdpy-30:RFP:NCAP-1:unc-54(3'UTR)] IV; apm-2(ox546[W64X]) mewSi79[Papm-2::3xFLAG:APM-2(T160A):tev-site unc-119(+)] X |
| strain, strain background (*C. elegans* , hermaphrodite) | GUN297 | This paper | | fcho-1(ox477::unc-119(+)) oxSi883[Phsp-16.41::TEVprotease unc-119(+)] ncap-1(mew39[1.4 kb deletion]) II; mewSi80[Pdpy-30:RFP:NCAP-1(K148A, E149A, G150A)] IV; apm-2(ox546[W64X]) oxSi877[Papm-2::3xFLAG:APM-2:tev-site unc-119(+)] X |
| strain, strain background (*C. elegans* , hermaphrodite) | GUN298 | This paper | | fcho-1(mew55) ncap-1(mew158[ncap-1(S84N):mScarlet:3xFLAG]) II |
| strain, strain background (*C. elegans* , hermaphrodite) | GUN299 | This paper | | fcho-1(mew55) ncap-1(mew54[ncap-1:mScarlet:3xFLAG]) II apa-2(mew159[V469D]) X |
| strain, strain background (*C. elegans* , hermaphrodite) | GUN300 | This paper | | fcho-1(mew55) ncap-1(mew54[ncap-1:mScarlet:3xFLAG]) II apb-1(mew160[S444P]) III |
| strain, strain background (*C. elegans* , hermaphrodite) | GUN301 | This paper | | fcho-1(mew55) ncap-1(mew54[ncap-1:mScarlet:3xFLAG]) II apb-1(mew161[S444P]) III |
| strain, strain background (*C. elegans* , hermaphrodite) | GUN302 | This paper | | fcho-1(mew55) ncap-1(mew54[ncap-1:mScarlet:3xFLAG]) II apb-1(mew162[S444P]) III |
| strain, strain background (*C. elegans* , hermaphrodite) | GUN303 | This paper | | fcho-1(mew55) ncap-1(mew54[ncap-1:mScarlet:3xFLAG]) II apm-2(mew163[T160A]) X |
| strain, strain background (*C. elegans* , hermaphrodite) | GUN304 | This paper | | fcho-1(mew55) ncap-1(mew54[ncap-1:mScarlet:3xFLAG]) II apm-2(mew164[T160A]) X |
| strain, strain background (*C. elegans* , hermaphrodite) | GUN305 | This paper | | fcho-1(mew55) ncap-1(mew54[ncap-1:mScarlet:3xFLAG]) II apm-2(mew165[T160A]) X |
| strain, strain background (*C. elegans* , hermaphrodite) | GUN306 | This paper | | fcho-1(mew55) ncap-1(mew54[ncap-1:mScarlet:3xFLAG]) II apm-2(mew166[T160A]) X |
| strain, strain background (*C. elegans* , hermaphrodite) | GUN307 | This paper | | mewSi81[Pdpy-30:apb-1(trunk):GFP:unc-54(3'UTR)] I; fcho-1(ox477::unc-119(+)) mewSi2[Pdpy-30:RFP:NCAP-1:unc-54(3'UTR)] ncap-1(mew39[1.4 kb deletion]) apm-2(mew168[T160A]) X |
| strain, strain background (*C. elegans* , hermaphrodite) | GUN308 | This paper | | mewSi81[Pdpy-30:apb-1(trunk):GFP:unc-54(3'UTR)] I; fcho-1(ox477::unc-119(+)) mewSi2[Pdpy-30:RFP:NCAP-1:unc-54(3'UTR)] ncap-1(mew39[1.4 kb deletion]) II apm-2(mew169[T160I]) X |
| strain, strain background (*C. elegans* , hermaphrodite) | GUN309 | This paper | | mewSi81[Pdpy-30:apb-1(trunk):GFP:unc-54(3'UTR)] I; fcho-1(ox477::unc-119(+)) mewSi2[Pdpy-30:RFP:NCAP-1:unc-54(3'UTR)] ncap-1(mew39[1.4 kb deletion]) II apm-2(mew170[T160I]) X |
| strain, strain background (*C. elegans* , hermaphrodite) | GUN343 | This paper | | fcho-1(ox477::unc-119(+)) oxSi883[Phsp-16.41::TEVprotease unc-119(+)] ncap-1(mew39[1.4 kb deletion]) II; mewSi3[Pdpy-30:RFP:NCAP-1:unc-54(3'UTR)] IV; apa-2(mew191[V469D]) apm-2(ox546[W64X]) oxSi877[Papm-2::3xFLAG:APM-2:tev-site unc-119(+)] X |
| strain, strain background (*C. elegans* , hermaphrodite) | GUN344 | This paper | | fcho-1(ox477::unc-119(+)) oxSi883[Phsp-16.41::TEVprotease unc-119(+)] ncap-1(mew39[1.4 kb deletion]) II; apb-1(mew192[S444P]) III; mewSi3[Pdpy-30:RFP:NCAP-1:unc-54(3'UTR)] IV; apm-2(ox546[W64X]) oxSi877[Papm-2::3xFLAG:APM-2:tev-site unc-119(+)] X |
| strain, strain background (*C. elegans* , hermaphrodite) | EG6703 | DOI: 10.1038/nmeth.1865 | | unc-119(ed3)III; cxTi10816 IV; oxEx1582[Peft-3::GFP unc-119(+)] |
| genetic reagent (*C. elegans* ) | ncap-1(mew54[C-terminal mScarlet:3xFlag]) | This paper | | Generated by CRISPR |
| genetic reagent (*C. elegans* ) | apa-2(mew126[V469D]) | This paper | | Generated by CRISPR |
| genetic reagent (*C. elegans* ) | apb-1(mew138[S444P]) | This paper | | Generated by CRISPR |
| genetic reagent (*C. elegans* ) | apb-1(mew150[S444P]) | This paper | | Generated by CRISPR |
| genetic reagent (*C. elegans* ) | apa-2(mew153[V469D]) | This paper | | Generated by CRISPR |
| genetic reagent (*C. elegans* ) | apm-2(mew156[T160A]) | This paper | | Generated by mutagenesis |
| genetic reagent (*C. elegans* ) | ncap-1(mew158[S84N, C-terminal mScarlet:3xFlag]) | This paper | | Generated by CRISPR |
| genetic reagent (*C. elegans* ) | apa-2(mew159[V469D]) | This paper | | Generated by mutagenesis |
| genetic reagent (*C. elegans* ) | apb-1(mew160[S444P]) | This paper | | Generated by mutagenesis |
| genetic reagent (*C. elegans* ) | apb-1(mew161[S444P]) | This paper | | Generated by mutagenesis |
| genetic reagent (*C. elegans* ) | apb-1(mew162[S444P]) | This paper | | Generated by mutagenesis |
| genetic reagent (*C. elegans* ) | apm-2(mew163[T160A]) | This paper | | Generated by mutagenesis |
| genetic reagent (*C. elegans* ) | apm-2(mew164[T160A]) | This paper | | Generated by mutagenesis |
| genetic reagent (*C. elegans* ) | apm-2(mew165[T160A]) | This paper | | Generated by mutagenesis |
| genetic reagent (*C. elegans* ) | apm-2(mew166[T160A]) | This paper | | Generated by mutagenesis |
| genetic reagent (*C. elegans* ) | apm-2(mew167[T160A]) | This paper | | Generated by mutagenesis |
| genetic reagent (*C. elegans* ) | apm-2(mew168[T160A]) | This paper | | Generated by mutagenesis |
| genetic reagent (*C. elegans* ) | apm-2(mew169[T160A]) | This paper | | Generated by mutagenesis |
| genetic reagent (*C. elegans* ) | apm-2(mew170[T160A]) | This paper | | Generated by mutagenesis |
| genetic reagent (*C. elegans* ) | apa-2(mew191[V469D]) | This paper | | Generated by CRISPR |
| genetic reagent (*C. elegans* ) | apb-1(mew192[S444P]) | This paper | | Generated by CRISPR |
| genetic reagent (*C. elegans* ) | mewSI32(Pdpy-30::TagRFP-T:ncap-1(S84N minigene)::unc-54UTR Cb_unc-119(+) at ttTi5605) II | This paper | | Generated by CRISPR |
| genetic reagent (*C. elegans* ) | mewSI35(Pdpy-30::TagRFP-T:ncap-1(S84N)::unc-54UTR Cb_unc-119(+) at ttTi10816) IV | This paper | | Generated by CRISPR |
| genetic reagent (*C. elegans* ) | mewSI38(Pdpy-30::TagRFP-T:ncap-1(130-236)::unc-54UTR Cb_unc-119(+) at ttTi10816) IV | This paper | | Generated by CRISPR |

| | | | | |
|---|---|---|---|---|
| genetic reagent (*C. elegans* ) | *mewSI40(Pdpy-30::TagRFP-T::unc-54UTR Cb_unc-119(+) at ttTi10816) IV* | This paper | | Generated by CRISPR |
| genetic reagent (*C. elegans* ) | *mewSI46(Pdpy-30::TagRFP-T::unc-54UTR Cb_unc-119(+) at ttTi5605) II* | This paper | | Generated by CRISPR |
| genetic reagent (*C. elegans* ) | *mewSI62(Pdpy-30::TagRFP-T:ncap-1(R109E,minigene)::unc-54UTR Cb_unc-119(+) at ttTi5605) II* | This paper | | Generated by CRISPR |
| genetic reagent (*C. elegans* ) | *mewSI65(Pdpy-30::TagRFP-T:ncap-1(R109E)::unc-54UTR Cb_unc-119(+) at ttTi10816)IV* | This paper | | Generated by CRISPR |
| genetic reagent (*C. elegans* ) | *mewSI66(Pdpy-30::TagRFP-T:ncap-1(K148A,E149A,G150A,minigene)::unc-54UTR Cb_unc-119(+) at ttTi5605) II* | This paper | | Generated by CRISPR |
| genetic reagent (*C. elegans* ) | *mewSI75(Pdpy-30::TagRFP-T:ncap-1(minigene)::unc-54UTR Cb_unc-119(+) at ttTi5605) II* | This paper | | Generated by CRISPR |
| genetic reagent (*C. elegans* ) | *mewSi79(Flag - apm-2(T160A) with TEV site (Papm-2), unc-119 rescue) X* | This paper | | Generated by CRISPR |
| genetic reagent (*C. elegans* ) | *mewSI80(Pdpy-30::TagRFP-T:ncap-1(K148A, E149A, G150A)::unc-54UTR Cb_unc-119(+) at ttTi10816) IV* | This paper | | Generated by CRISPR |
| genetic reagent (*C. elegans* ) | *mewSi81(Pdpy-30::apb-1(trunk):GFP::unc-54UTR Cb_unc-119(+) at ttTi4348) I* | This paper | | Generated with MosSCI |
| genetic reagent (*C. elegans* ) | *mewSi82(Pdpy-30::apb-1(trunk):GFP::unc-54UTR Cb_unc-119(+) at ttTi4348) I* | This paper | | Generated by CRISPR |
| antibody | rabbit anti-AP2M1 phospho T156 | Abcam | Cat# 109397, RRID:AB_10866362 | (1:1000) |
| antibody | goat anti-rabbit Alexa Fluor 647 | Life Technologies | Cat# A21244, RRID: AB_1562581 | (1:2000) |
| antibody | mouse anti-flag | Sigma-Aldrich | Cat# F3165, RRID: AB_259529 | (1:1000) |
| antibody | goat anti-mouse IRDye 800CW | LI-COR | Cat# 925-32210, RRID: AB_2687825 | (1:20000) |
| recombinant DNA reagent | pEP1 | This paper | | See Supplementary File 1 |
| recombinant DNA reagent | pEP57 | This paper | | See Supplementary File 1 |
| recombinant DNA reagent | pEP82 | DOI: 10.7554/eLife.32242.001 | | |
| recombinant DNA reagent | pEP213 | This paper | | See Supplementary File 1 |
| recombinant DNA reagent | pEP218 | This paper | | See Supplementary File 1 |
| recombinant DNA reagent | pEP220 | This paper | | See Supplementary File 1 |
| recombinant DNA reagent | pEP221 | This paper | | See Supplementary File 1 |
| recombinant DNA reagent | pEP223 | This paper | | See Supplementary File 1 |
| recombinant DNA reagent | pEP239 | This paper | | See Supplementary File 1 |
| recombinant DNA reagent | pEP241 | This paper | | See Supplementary File 1 |
| recombinant DNA reagent | pEP242 | This paper | | See Supplementary File 1 |
| recombinant DNA reagent | pEP243 | This paper | | See Supplementary File 1 |
| recombinant DNA reagent | pEP244 | This paper | | See Supplementary File 1 |
| recombinant DNA reagent | pEP245 | This paper | | See Supplementary File 1 |
| recombinant DNA reagent | pEP246 | This paper | | See Supplementary File 1 |
| recombinant DNA reagent | pGB28 | DOI: 10.7554/eLife.32242.001 | | |
| recombinant DNA reagent | pGB51 | This paper | | See Supplementary File 1 |
| recombinant DNA reagent | pGB63 | This paper | | See Supplementary File 1 |
| recombinant DNA reagent | pGB73 | This paper | | See Supplementary File 1 |
| recombinant DNA reagent | pGB76 | This paper | | See Supplementary File 1 |
| recombinant DNA reagent | pGB86 | This paper | | See Supplementary File 1 |
| recombinant DNA reagent | pGB103 | This paper | | See Supplementary File 1 |
| recombinant DNA reagent | pGB104 | This paper | | See Supplementary File 1 |
| recombinant DNA reagent | pGB106 | This paper | | See Supplementary File 1 |
| recombinant DNA reagent | pGH419 | This paper | | See Supplementary File 1 |
| recombinant DNA reagent | pGH500 | DOI: 10.7554/eLife.32242.001 | | |
| recombinant DNA reagent | pGH503 | DOI: 10.7554/eLife.32242.001 | | |
| recombinant DNA reagent | pGH504 | DOI: 10.7554/eLife.03648.001 | | |
| recombinant DNA reagent | pSEM87 | DOI: 10.1534/g3.117.040824 | | |
| sequence-based reagent | oEP318 | Integrated DNA Technologies | | CATCATCACCATCACCACTAA |
| sequence-based reagent | oEP324 | Integrated DNA Technologies | | ATGTATATCTCCTTCTTAAAGTTAAACAAAATTAT |
| sequence-based reagent | oEP366 | Integrated DNA Technologies | | AACGGGCGGTAGTGGAGGCACTGGTATGGGAGATTACGAGAACGTTTTAATG |
| sequence-based reagent | oEP369 | Integrated DNA Technologies | | TATCACCACTTTGTACAAGAAAGCTGGGTCTAACTTTTATCCTTTTTTCCAATGTTAATT |
| sequence-based reagent | oEP512 | Integrated DNA Technologies | | AGTACTAGCGGTGGCAGTGGAGGTACCGGCGGAAGCATGGTCAGCAAGGGAGAGGCAGTT |
| sequence-based reagent | oEP513 | Integrated DNA Technologies | | CCGTCTTTATAGTCACCATCGTGGTCTTTGTAGTCCTTGTAGAGCTCGTCCATTCCTCCG |
| sequence-based reagent | oEP519 | Integrated DNA Technologies | | TTCCGACTAAAAATCCCCAAATTTTCAGAGATTTCAGTACTAGCGGTGGCAGTGGAGGTA |
| sequence-based reagent | oEP520 | Integrated DNA Technologies | | AGTTGTACGGAGAAGAAAGATGACGTCATCGCTTATCCTCCTTTGTCGTCATCATCCTTA |
| sequence-based reagent | oEP582 | Integrated DNA Technologies | | GGAGGAACGGGCGGTAGTGGAGGCACTGGTTAGACCTAGCTTTCTTGTACAAAGTGGTGA |
| sequence-based reagent | oEP584 | Integrated DNA Technologies | | GGAGGAACGGGCGGTAGTGGAGGCACTGGTGCGGAACTGGAAAAACAGGATCTTTCTGCC |
| sequence-based reagent | oEP641 | Integrated DNA Technologies | | GCCTTCTTTTTTTTTCATGTTGG |
| sequence-based reagent | oEP655 | Integrated DNA Technologies | | AAAAAGTCGATAGAGAAGGCTTCAACACAC |
| sequence-based reagent | oEP656 | Integrated DNA Technologies | | TCGGTCAAATTTCCCGGTTTTTAAC |
| sequence-based reagent | oEP657 | Integrated DNA Technologies | | ACCCGATTTTCTCGGTTTTTCTCTC |
| sequence-based reagent | oEP659 | Integrated DNA Technologies | | ATAAAAGTACGGATTTTTGCTCGAAATCAAC |
| sequence-based reagent | oEP660 | Integrated DNA Technologies | | GGCCAATTTTGAGGATTCTTTGGC |
| sequence-based reagent | oEP661 | Integrated DNA Technologies | | TTTAGACTGAAAATTCCGATTTTTGAGCC |
| sequence-based reagent | oEP662 | Integrated DNA Technologies | | GGTGGAGAGAGAGAAGTGAAGAGACGC |
| sequence-based reagent | oEP670 | Integrated DNA Technologies | | GCAAACTGGGGCACAAACTTAATTCC |
| sequence-based reagent | oEP793 | Integrated DNA Technologies | | GATCACTTTCGTTATATCGAACGAAGCTAATTCTTGTACAAAGTGGTGATATCTGAGCTC |
| sequence-based reagent | oEP808 | Integrated DNA Technologies | | AACAACATGAAGTGGCAGTCGC |
| sequence-based reagent | oEP812 | Integrated DNA Technologies | | GGAGCACAGGGAGAAAGAGC |
| sequence-based reagent | oEP894 | Integrated DNA Technologies | | AGGCATTCGTGGGATGCGGGTTTCAAGAAGAGGGAGATGCTTTTGACTTTAATGTCACAC |
| sequence-based reagent | oEP865 | Integrated DNA Technologies | | GGATGCGGGTTTCAAGAAGAG |
| sequence-based reagent | oEP969 | Integrated DNA Technologies | | CGTCAAGTGAAGCTGATCCGTGGCGTAGTTGGCTCACTTGGATTAATCGTAATCCC |

| | | | | |
|---|---|---|---|---|
| sequence-based reagent | oEP970 | Integrated DNA Technologies | | GGGATTACGATTAATCCAAGTGAGCCAACTACGCCACGGATCAGCTTCACTTGACG |
| sequence-based reagent | oEP971 | Integrated DNA Technologies | | CGTCAAGTGAAGCTGATCCGTGGCGTAGTTGGCTCACTTGGATTAATCGTAATCCCGCTA |
| sequence-based reagent | oEP97 | Integrated DNA Technologies | | TAGCGGGATTACGATTAATCCAAGTGAGCCAACTACGCCACGGATCAGCTTCACTTGACG |
| sequence-based reagent | oEP973 | Integrated DNA Technologies | | TGCTCGAGCACCTATGATTTGGATTGTAGGAGAATATGC |
| sequence-based reagent | oEP974 | Integrated DNA Technologies | | TAGGTGCTCGAGCATCGGGTTCATCCAG |
| sequence-based reagent | oEP977 | Integrated DNA Technologies | | ATTGTCGACAATCGCGATGATGTGCAG |
| sequence-based reagent | oEP978 | Integrated DNA Technologies | | CGATTGTCGACAATCTGGATGACGCG |
| sequence-based reagent | oEP985 | Integrated DNA Technologies | | TGCCGGTCCAAGCTTGGATTTAGCATTTGCAGCTGCACAAACCATCTCAATTAACATTGG |
| sequence-based reagent | oEP987 | Integrated DNA Technologies | | GTTATTATTTCGGATTGTCGACAATAGGGAGGACGTTCAAGGATACGCAGCAAAGACTGT |
| sequence-based reagent | oEP1010 | Integrated DNA Technologies | | ACCGATAACAGCCGTTATTTTGTTATTCGTA |
| sequence-based reagent | oEP1011 | Integrated DNA Technologies | | ACGGCTGTTATCGGTCACGCT |
| sequence-based reagent | oEP1014 | Integrated DNA Technologies | | GGGACAAACCATCTCAATTAACATTGGAAAAAAAGACAAGTCATAGACTCAGCTTTCTTGTACAAAGTGGTGATATCTGA |
| sequence-based reagent | oEP1016 | Integrated DNA Technologies | | CAGTGAAAAGTTCTTCTCCTTTACT |
| sequence-based reagent | oEP1020 | Integrated DNA Technologies | | TAGTGCCACTGCTTCCACCGCCGCCAGGTGCCGGCTAAACCCAGCTTTCTTGTACAAAGT |
| sequence-based reagent | oEP1031 | Integrated DNA Technologies | | CATCACCATCATCACCACTGAGATC |
| sequence-based reagent | oEP1032 | Integrated DNA Technologies | | CATGCTTCCGCCGGTAC |
| sequence-based reagent | oEP1033 | Integrated DNA Technologies | | AGGTACCGGCGGAAGCATGGAAGAAAGCGGCTATG |
| sequence-based reagent | oEP1034 | Integrated DNA Technologies | | TCAGTGGTGATGATGGTGATGGCCTTCTTTTTTTTTCATGTTGGC |
| sequence-based reagent | oEP1035 | Integrated DNA Technologies | | TCAGTGGTGATGATGGTGATGCTGCTGTTTAACCCATTTAAAGTG |
| sequence-based reagent | oEP103 | Integrated DNA Technologies | | GGTACCGGCGGAAGCTGCGAATTTGCAAAACAGGC |
| sequence-based reagent | oEP1037 | Integrated DNA Technologies | | TGATGAGGGTGATGCCTTTGATTTTAATG |
| sequence-based reagent | oEP1038 | Integrated DNA Technologies | | CACCCTCATCACCAAAACCAATACCAAT |
| sequence-based reagent | oEP1041 | Integrated DNA Technologies | | TTTTGCAGCAGCTCAGACCATCAAACTGAACATTG |
| sequence-based reagent | oEP1042 | Integrated DNA Technologies | | CTGAGCTGCTGCAAAACCCAGATCCAGTTTCGG |
| sequence-based reagent | oEP1051 | Integrated DNA Technologies | | TCAGTGGTGATGATGGTGATGTTTCCCACCCGGTGGTG |
| sequence-based reagent | oGB24 | Integrated DNA Technologies | | CGCCGCCAGCCAATCTGCCCAGCCACCTGGCTGGTGATCTGGGACTGTTC |
| sequence-based reagent | oGB26 | Integrated DNA Technologies | | ATGAATAAGCCTCCGATCATCATATGTATATCTCCTTCTTATA |
| sequence-based reagent | oGB27 | Integrated DNA Technologies | | GCATTTATGAAACCCGCTGCTAATTAACCTAGGCTGCTGCCACCG |
| sequence-based reagent | oGB28 | Integrated DNA Technologies | | ATGATCGGAGGCTTATTCATCT |
| sequence-based reagent | oGB29 | Integrated DNA Technologies | | GCAGCGGGTTTCATAAATGCCA |
| sequence-based reagent | oGB33 | Integrated DNA Technologies | | GGGCAGATTGGCTGGCGGCGAGAAGGCATCAAGTA |
| sequence-based reagent | oGB34 | Integrated DNA Technologies | | AGCAAGAGTCTGGTGCCGCGCGGCAGCGGA |
| sequence-based reagent | oGB35 | Integrated DNA Technologies | | CTGCTTACCGCTGCCGCGCGGCACCAGACCTTGCTTGTTTCATCAGCTGTG |
| sequence-based reagent | oGB52 | Integrated DNA Technologies | | TGCATCACGGGAGATGCACT |
| sequence-based reagent | oGB124 | Integrated DNA Technologies | | GGCTGGGTCCAGTTCCATCACCATCATCACCACTGA |
| sequence-based reagent | oGB125 | Integrated DNA Technologies | | GAACTGGACCCAGCCGGTGC |
| sequence-based reagent | oGB130 | Integrated DNA Technologies | | GGAGCAGTCACAAATCACGTCTCAAGTTGCCGGCCAAATTGGATGGCGTCGGGAGGGTAT |
| sequence-based reagent | oGB147 | Integrated DNA Technologies | | ATCTCCCGTGATGCAGGGCCTGGCTCTTGGGGTAC |
| sequence-based reagent | oGB148 | Integrated DNA Technologies | | TGTGAGTTTGCGAAACAAGC |
| sequence-based reagent | oGB149 | Integrated DNA Technologies | | TTTCGCAAACTCACACATGCTTCCGCCGGTACCTC |
| sequence-based reagent | oGB172 | Integrated DNA Technologies | | ATGATCGGAGGCTTATTCATCT |
| sequence-based reagent | oGB173 | Integrated DNA Technologies | | TAGATGAATAAGCCTCCGATCATATGTATATCTCCTTCT |
| sequence-based reagent | oGB175 | Integrated DNA Technologies | | CATCACCATCATCACCAC |
| sequence-based reagent | oGB180 | Integrated DNA Technologies | | CAGTGGTGATGATGGTGATGGCTTCCGCCGGTACCTCCAC |
| sequence-based reagent | oGH367 | Integrated DNA Technologies | | TAATGCTTAAGTCGAACAGAAAGTAATCG |
| sequence-based reagent | oGH369 | Integrated DNA Technologies | | TCTGTTCGACTTAAGCATTATTTGCGATGAATCCCATGAC |
| sequence-based reagent | oGH408 | Integrated DNA Technologies | | GCATTTTTCACATTTTCTAACATTTTTTCTGTTGAAAAG |
| sequence-based reagent | oGH409 | Integrated DNA Technologies | | GCACATTTTAAGTCTGTAAAAGTGAAAACCCA |
| sequence-based reagent | oGH411 | Integrated DNA Technologies | | TCCTATGCTCAGTCAGTGTATGAGC |
| sequence-based reagent | oGH412 | Integrated DNA Technologies | | GCTTTTGGAGCATTTTGTTTTCTAATTTTGAATGA |
| sequence-based reagent | oGH413 | Integrated DNA Technologies | | AAGTTTTATACCAAGTTTAGAACATGGATTCGG |
| sequence-based reagent | oGH414 | Integrated DNA Technologies | | CGGTTCTGCATGCAGTTGTCTG |
| sequence-based reagent | oGH415 | Integrated DNA Technologies | | CCAAAAAAATGTATCTGAATAAGTAAAGCAAAGTGATTC |
| sequence-based reagent | oGH416 | Integrated DNA Technologies | | GTCTTAACCAAAGAGCAACAACAATACCT |
| sequence-based reagent | oGH417 | Integrated DNA Technologies | | ACTATAACTTTTGATTGTTTTGTCAACAGCTAGC |
| sequence-based reagent | oGH418 | Integrated DNA Technologies | | CCACTTTTTCTAATATTTCAAACTTGTGCTCGA |
| sequence-based reagent | oGH419 | Integrated DNA Technologies | | TGTAAAAGTGGAGAGATGGCACGG |
| sequence-based reagent | oGH430 | Integrated DNA Technologies | | GTTTTGCAAAGATATTTTAATGAAGTTTGGCTCA |
| sequence-based reagent | oGH432 | Integrated DNA Technologies | | AAATTAATTGTTTCTACAGAGTGTTTCAATGTTTGAAC |
| sequence-based reagent | oGH441 | Integrated DNA Technologies | | GCTCCAATTTCCTTGAAACCTCG |
| sequence-based reagent | oGH442 | Integrated DNA Technologies | | CCTTGAAAGCTTTTTTTAAGTTTTTTAGGTG |
| sequence-based reagent | oGH443 | Integrated DNA Technologies | | GATTTTTCAAAATTTTTAACATCGAAACTCCC |
| sequence-based reagent | oGH444 | Integrated DNA Technologies | | GCCCGATTTTACAGGAACTCC |
| sequence-based reagent | oGH445 | Integrated DNA Technologies | | CTAAAATTCTAAACTACAAAATAATAATAAAAATATC |
| sequence-based reagent | oGH446 | Integrated DNA Technologies | | TGCAATTTTTACAGGTCAGG |
| sequence-based reagent | oGH447 | Integrated DNA Technologies | | CTCGGAAATTCAAATTATACATCAAAAATTATCAC |
| sequence-based reagent | oGH448 | Integrated DNA Technologies | | GAAATTCAGAATTATTTAGGGGAAAAGGC |
| sequence-based reagent | oGH452 | Integrated DNA Technologies | | CCATTCATATTTTGTCTCAGGAGAATAC |
| sequence-based reagent | oGH679 | Integrated DNA Technologies | | AGGTATTCAGACATTTTTCAAATGAAAATCTAC |

| | | | | |
|---|---|---|---|---|
| sequence-based reagent | oGH847 | Integrated DNA Technologies | | CCAAACTGAAGGTCAAGGTGGTC |
| sequence-based reagent | oGH848 | Integrated DNA Technologies | | CCTTGACCTTCAGTTTGGTGCGC |
| sequence-based reagent | oGH1014 | Integrated DNA Technologies | | CCGCCGTCGTTCTCTCCACCG |
| sequence-based reagent | RWB099 | Integrated DNA Technologies | | GGCCTCCTTCGTCGTCTTCAGGATCCAATTCGA GCTCGAACAACAAC |
| sequence-based reagent | rEP578 | Integrated DNA Technologies | | CGGTAGTGGAGGCACTGGTA |
| sequence-based reagent | rEP579 | Integrated DNA Technologies | | ACCACTTTGTACAAGAAAGC |
| sequence-based reagent | rEP580 | Integrated DNA Technologies | | AAGATCCTGTTTTTCCAGTT |
| sequence-based reagent | rEP807 | Integrated DNA Technologies | | GGATGCGGGTTTCAGGAGAG |
| sequence-based reagent | rEP905 | Integrated DNA Technologies | | CTTGGATTTAGCATTTAAAG |
| sequence-based reagent | rEP980 | Integrated DNA Technologies | | TAATCCAAATCATTGAAGCC |
| sequence-based reagent | rEP986 | Integrated DNA Technologies | | CGTATCCTTGAACGTCCTCT |
| sequence-based reagent | rEP1013 | Integrated DNA Technologies | | CTCAATTAACATTGGAAAAA |
| sequence-based reagent | rEP1019 | Integrated DNA Technologies | | CTGATCATTGAGCCGGCACC |
| sequence-based reagent | rGB156 | Integrated DNA Technologies | | CAAATCACGTCTCAAGTGAC |
| sequence-based reagent | rKW3 | Integrated DNA Technologies | | CTTAGAAATCTCTGAAAATT |
| peptide, recombinant protein | AcTEV Protease | Invitrogen | 12575015 | |
| peptide, recombinant protein | Thrombin Protease | Sigma | T7009 | |
| Chemical compound, drug | Inositol Hexakisphosphate (Phytic Acid or IP6) | Sigma | P9910 | |
| Chemical compound, drug | Heparin | Sigma | H3393 | |
| software, algorithm | GraphPad Prism (version 7 for Windows) | GraphPad Software, www.graphpad.com | RRID:SCR_002798 | |
| software, algorithm | Fiji | doi:10.1038/nmeth.2019 | RRID:SCR_002285 | |
