## Supplemental File 1 for "A structural mechanism for phosphorylation-dependent inactivation of the AP2 complex"

**This supplementary file contains the following information associated with '*A structural mechanism for phosphorylation-dependent inactivation of the AP2 complex*'**

*C. elegans* Strains

Recombinant proteins

DNA plasmids (including cloning strategies)

Sequences of synthesized gene fragments (IDT gBlocks)

Sequences of DNA oligos

***C. elegans* Strains**

### Wild-Type

Bristol N2 *wild-type*

### Genetic screen parent strains

GUN61 *mewSi1[Pdpy-30:apb-1(trunk):GFP:unc-54(3'UTR)] I; fcho-1(ox477::unc-119(+)) mewSi2[Pdpy-30:RFP:NCAP-1:unc-54(3'UTR)] ncap-1(mew39[1.4 kb deletion]) II*

GUN141 *fcho-1(mew55) ncap-1(mew54[ncap-1:mScarlet:3xFlag]) II*

### Genetic screen results

GUN294 *fcho-1(mew55) ncap-1(mew54[ncap-1:mScarlet:3xFLAG]) II apm-2(mew156[T160A]) X*

GUN298 *fcho-1(mew55) ncap-1(mew158[ncap-1(S84N):mScarlet:3xFLAG]) II*

GUN299 *fcho-1(mew55) ncap-1(mew54[ncap-1:mScarlet:3xFLAG]) II apa-2(mew159[V469D]) X*

GUN300 *fcho-1(mew55) ncap-1(mew54[ncap-1:mScarlet:3xFLAG]) II apb-1(mew160[S444P]) III*

GUN301 *fcho-1(mew55) ncap-1(mew54[ncap-1:mScarlet:3xFLAG]) II apb-1(mew161[S444P]) III*

GUN302 *fcho-1(mew55) ncap-1(mew54[ncap-1:mScarlet:3xFLAG]) II apb-1(mew162[S444P]) III*

GUN303 *fcho-1(mew55) ncap-1(mew54[ncap-1:mScarlet:3xFLAG]) II apm-2(mew163[T160A]) X*

GUN304 *fcho-1(mew55) ncap-1(mew54[ncap-1:mScarlet:3xFLAG]) II apm-2(mew164[T160A]) X*

GUN305 *fcho-1(mew55) ncap-1(mew54[ncap-1:mScarlet:3xFLAG]) II apm-2(mew165[T160A]) X*

GUN306 *fcho-1(mew55) ncap-1(mew54[ncap-1:mScarlet:3xFLAG]) II apm-2(mew166[T160A]) X*

GUN307 *mewSi81[Pdpy-30:apb-1(trunk):GFP:unc-54(3'UTR)] I; fcho-1(ox477::unc-119(+)) mewSi2[Pdpy-30:RFP:NCAP-1:unc-54(3'UTR)] ncap-1(mew39[1.4 kb deletion]) apm-2(mew168[T160A]) X*

GUN308 *mewSi81[Pdpy-30:apb-1(trunk):GFP:unc-54(3'UTR)] I; fcho-1(ox477::unc-119(+)) mewSi2[Pdpy-30:RFP:NCAP-1:unc-54(3'UTR)] ncap-1(mew39[1.4 kb deletion]) II apm-2(mew169[T160I]) X*

GUN309 *mewSi81[Pdpy-30:apb-1(trunk):GFP:unc-54(3'UTR)] I; fcho-1(ox477::unc-119(+)) mewSi2[Pdpy-30:RFP:NCAP-1:unc-54(3'UTR)] ncap-1(mew39[1.4 kb deletion]) II apm-2(mew170[T160I]) X*

**<u>Fitness assay and *in vivo* protease assay</u>**

*PHear Interface (Figures 2C and 2F)*

GUN198 *fcho-1(ox477::unc-119(+)) oxSi883[Phsp-16.41::TEVprotease unc-119(+)] ncap-1(mew39[1.4 kb deletion]) II; mewSi40[Pdpy-30:RFP:unc-54(3'UTR)] IV; apm-2(ox546[W64X]) oxSi877[Papm-2::3xFLAG:APM-2:tev-site unc-119(+)] X*

*Ex interface fitness assay (Figure 6C)*

GUN87 *ncap-1(mew54[ncap-1:mScarlet:3xFLAG]) II*

GUN88 *ncap-1(mew54[ncap-1:mScarlet:3xFLAG]) fcho-1(ox477::unc-119(+)) II*

GUN291 *ncap-1(mew54[ncap-1:mScarlet:3xFLAG]) fcho-1(ox477::unc-119(+)) II; apa-2(mew153[V469D]) X*

GUN288 *ncap-1(mew54[ncap-1:mScarlet:3xFLAG]) fcho-1(ox477) II; apb-1(mew150[S444P]) III*

*NECAP structure-function (Figures 7A bottom and 7B)*

GUN198 *fcho-1(ox477::unc-119(+)) oxSi883[Phsp-16.41::TEVprotease unc-119(+)] ncap-1(mew39[1.4 kb deletion]) II; mewSi40[Pdpy-30:RFP:unc-54(3'UTR)] IV; apm-2(ox546[W64X]) oxSi877[Papm-2::3xFLAG:APM-2:tev-site unc-119(+)] X*

GUN106 *fcho-1(ox477::unc-119(+)) oxSi883[Phsp-16.41::TEVprotease unc-119(+)] ncap-1(mew39[1.4 kb deletion]) II; mewSi3[Pdpy-30:RFP:NCAP-1:unc-54(3'UTR)] IV; apm-2(ox546[W64X]) oxSi877[Papm-2::3xFLAG:APM-2:tev-site unc-119(+)] X*

GUN276 *fcho-1(ox477::unc-119(+)) oxSi883[Phsp-16.41::TEVprotease unc-119(+)] ncap-1(mew39[1.4 kb deletion]) II; mewSi75[Pdpy-30:RFP:NCAP-1(1-190)] IV; apm-2(ox546[W64X]) oxSi877[Papm-2::3xFLAG:APM-2:tev-site unc-119(+)] X*

GUN89 *fcho-1(ox477::unc-119(+)) oxSi883[Phsp-16.41::TEVprotease unc-119(+)] ncap-1(mew39[1.4 kb deletion]) II; mewSi12[Pdpy-30:RFP:NCAP-1(1-163)] IV; apm-2(ox546[W64X]) oxSi877[Papm-2::3xFLAG:APM-2:tev-site unc-119(+)] X*

GUN195 *fcho-1(ox477::unc-119(+)) oxSi883[Phsp-16.41::TEVprotease unc-119(+)] ncap-1(mew39[1.4 kb deletion]) II; mewSi37[Pdpy-30:RFP:NCAP-1(1-129)] IV; apm-2(ox546[W64X]) oxSi877[Papm-2::3xFLAG:APM-2:tev-site unc-119(+)] X*

GUN196 *fcho-1(ox477::unc-119(+)) oxSi883[Phsp-16.41::TEVprotease unc-119(+)] ncap-1(mew39[1.4 kb deletion]) II; mewSi38[Pdpy-30:RFP:NCAP-1(130-236)] IV; apm-2(ox546[W64X]) oxSi877[Papm-2::3xFLAG:APM-2:tev-site unc-119(+)] X*

### ***In vivo* imaging assay**

*PHear interface (Figure 3E)*

GUN204 *mewSi81[Pdpy-30:apb-1(trunk):GFP:unc-54(3'UTR)] I; fcho-1(ox477::unc-119(+)) mewSi46[Pdpy-30:RFP:unc-54(3'UTR)] ncap-1(mew39[1.4 kb deletion]) II; apm-2(ox562[E306K])X*

GUN62 *mewSi81[Pdpy-30:apb-1(trunk):GFP:unc-54(3'UTR)] I; fcho-1(ox477::unc-119(+)) mewSi2[Pdpy-30:RFP:NCAP-1:unc-54(3'UTR)] ncap-1(mew39[1.4 kb deletion]) II; apm-2(ox562[E306K])X*

GUN55 *mewSi81[Pdpy-30:apb-1(trunk):GFP:unc-54(3'UTR)] I; fcho-1(ox477::unc-119(+)) mewSi2[Pdpy-30:RFP:NCAP-1:unc-54(3'UTR)] ncap-1(mew39[1.4 kb deletion]) II; apm-2(ox562[E302K]+mew46[T160A])X*

GUN123 *mewSi81[Pdpy-30:apb-1(trunk):GFP:unc-54(3'UTR)] I; fcho-1(ox477::unc-119(+)) mewSi32[Pdpy-30:RFP:NCAP-1:unc-54(3'UTR)(S84N)] ncap-1(mew39[1.4 kb deletion]) II; apm-2(ox562[E306K])X*

GUN249 *mewSi81[Pdpy-30:apb-1(trunk):GFP:unc-54(3'UTR)] I; fcho-1(ox477::unc-119(+)) mewSi62[Pdpy-30:RFP:NCAP-1:unc-54(3'UTR)(R109E)] ncap-1(mew39[1.4 kb deletion]) II; apm-2(ox562[E306K])X*

*Ex Interface (Figure 6D)*

GUN204 *mewSi81[Pdpy-30:apb-1(trunk):GFP:unc-54(3'UTR)] I; fcho-1(ox477::unc-119(+)) mewSi46[Pdpy-30:RFP:unc-54(3'UTR)] ncap-1(mew39[1.4 kb deletion]) II; apm-2(ox562[E306K])X*

GUN62 *mewSi81[Pdpy-30:apb-1(trunk):GFP:unc-54(3'UTR)] I; fcho-1(ox477::unc-119(+)) mewSi2[Pdpy-30:RFP:NCAP-1:unc-54(3'UTR)] ncap-1(mew39[1.4 kb deletion]) II; apm-2(ox562[E306K])X*

GUN256 *mewSi81[Pdpy-30:apb-1(trunk):GFP:unc-54(3'UTR)] I; fcho-1(ox477::unc-119(+)) mewSi2[Pdpy-30:RFP:NCAP-1:unc-54(3'UTR)] ncap-1(mew39[1.4 kb deletion]) II; apa-2(mew126[V469D]) apm-2(ox562[E306K])X*

GUN271 *mewSi82[apb-1(trunk, S444P)] I; fcho-1(ox477::unc-119(+)) mewSi2[Pdpy-30:RFP:NCAP-1:unc-54(3'UTR)] ncap-1(mew39[1.4 kb deletion]) II; apb-1(mew138[S444P]) III; apm-2(ox562[E306K])X*

GUN253 *mewSi1[apb-2(trunk):GFP] I; fcho-1(ox477::unc-119(+)) mewSi66[Pdpy-30:RFP:NCAP-1:unc-54(3'UTR)(K148A,E149A,G150A)] ncap-1(mew39[1.4 kb deletion]) II; apm-2(ox562[E306K])X*

#### *Recombinant Proteins*

Recombinant protein sequences were derived from those below, with the addition of mutations, affinity tags, linkers, or other biochemical tools.

| | |
|---|---|
| Mouse AAK1 | XP_006506255 |
| Mouse AP2 beta1 | NP_001030931 |
| Mouse AP2 mu1 | NP_033809 |
| Mouse AP2 alpha2 | NP_001343997 |
| Rat AP2 sigma1 | NP_075241.2 |
| Human NECAP2 | NP_060560 |
| Mouse NECAP2 | NP_079659 |

#### *DNA Plasmids*

**NECAP expression plasmids**

*Controls (Figure 1C, 3C, 4F, 5A, 6B, 7A top)*

| | |
|---|---|
| pGB86 | *HaloTag(TEVsite):6xHis in pET-21b* |
| pEP223 | *HaloTag(TEVsite):human NECAP2(1-263):6xHis in pET-21b* |

*Structure Function (Figure 1C, 5A, 7A top)*

| | |
|---|---|
| pEP239 | *HaloTag(TEVsite):human NECAP2(1-195):6xHis in pET-21b* |
| pEP220 | *HaloTag(TEVsite):human NECAP2(1-170):6xHis in pET-21b* |
| pEP221 | *HaloTag(TEVsite):human NECAP2(1-132):6xHis in pET21b* |
| pEP241 | *HaloTag(TEVsite):human NECAP2(133-263):6xHis in pET21b* |

*PHear Interface Mutants (Figure 3C)*

pEP242 *HaloTag(TEVsite):human NECAP2(S87N, 1-263):6xHis in pET-21b*

pEP245 *HaloTag(TEVsite):human NECAP2(R112E, 1-263):6xHis in pET-21b*

*Ex Interface Mutants (Figure 6B)*

pEP243 *HaloTag(TEVsite):human NECAP2(K153A, E154A, G155A, 1-263):6xHis in pET-21b*

**AP2 expression plasmids**

*Phospho-AP2 (Figure 1C, 3C)*

pGH504 *mouse AP2 alpha2(1-621):GST + rat AP2 sigma1(1-142) in pACYCduet-1* (Hollopeter *et al*., 2014)

pGB104 *mouse AP2 beta1(1-591) + mouse AP2 mu1(1-435, thrombin site) in pETduet-1*

pEP82 *mouse AAK1(1-325) in pRSFduet* (Beacham *et al*., 2018)

*Phospho-site mutant (Figure 3C)*

pEP246 *mouse AP2 beta1(1-591) + mouse AP2 mu1(T156A, 1-435, thrombin site) in pETduet-1*

*AP2 muE302K (Figure 4F, 5A, 6B)*

pGH504 *mouse AP2 alpha2(1-621):GST + rat AP2 sigma1(1-142) in pACYCduet-1* (Hollopeter *et al.*, 2014)

pGB106 *mouse AP2 beta1(1-591) + mouse AP2 mu1(E302K, 1-435, thrombin site) in pETduet-1*

*Ex interface mutants (Figure 6B)*

pEP218 *mouse AP2 alpha2(I470D, 1-621):GST + rat AP2 sigma1(1-142) in pACYCduet-1*

pEP213 *mouse AP2 beta1(A447P, 1-591) + mouse AP2 mu1(E302K, 1-435, thrombin site) in pETduet-1*

*Phospho-AP2 muE302K (Figure 7A top)*

pGH504 *mouse AP2 alpha2(1-621):GST + rat AP2 sigma1(1-142) in pACYCduet-1* (Hollopeter *et al.*, 2014)

pGB106 *mouse AP2 beta1(1-591) + mouse AP2 mu1(E302K, 1-435, thrombin site) in pETduet-1*

pEP82 *mouse AAK1(1-325) in pRSFduet* (Beacham *et al.*, 2018)

**Plasmids for cryo-EM**

*NECAP*

pGH503 *Mouse NECAP2(1-266):6xHis in pET-21b* (Beacham *et al.*, 2018)

*Phospho-AP2 (unclamped, PDB 6OWO)*

pGH504 *mouse AP2 alpha2(1-621):GST + rat AP2 sigma1(1-142) in pACYCduet-1* (Hollopeter *et al*., 2014)

pGH419 *mouse AP2 beta1(1-591) + mouse AP2 mu1(1-435) in pETduet-1*

pEP82 *mouse AAK1(1-325) in pRSFduet* (Beacham *et al*., 2018)

*Closed AP2 (Figure 4D, 4E)*

pGH504 *mouse AP2 alpha2(1-621):GST + rat AP2 sigma1(1-142) in pACYCduet-1* (Hollopeter *et al*., 2014)

pGH419 *mouse AP2 beta1(1-591) + mouse AP2 mu1(1-435) in pETduet-1*

*Open AP2 (Figure 4D, 4E)*

pGH504 *mouse AP2 alpha2(1-621):GST + rat AP2 sigma1(1-142) in pACYCduet-1* (Hollopeter *et al*., 2014)

pGB103 *mouse AP2 beta1(1-591) + mouse AP2 mu1(E302K, 1-435) in pETduet-1*

*Phospho-AP2 (clamped, PDB 6OXL)*

pGH504 *mouse AP2 alpha2(1-621):GST + rat AP2 sigma1(1-142) in pACYCduet-1* (Hollopeter *et al*., 2014)

pGB103 *mouse AP2 beta1(1-591) + mouse AP2 mu1(E302K, 1-435) in pETduet-1*

pEP82 *mouse AAK1(1-325) in pRSFduet* (Beacham *et al*., 2018)

**<u>Other plasmids</u>**

*MosSCI plasmid to generate PHearEx strain (Figure 7A bottom, Figure 7B)*

pEP57 *Pdpy-30:tagRFP-T:NCAP-1(1-163):unc-54(3'UTR) (cxTi10816 targeting)*

*PCR template to generate ncap-1:mScarlet CRISPR repair template*

pSEM87 *twk-18:mScarlet CRISPR repair* (El Mouridi *et al*., 2017)

**Cloning of Plasmids**

The plasmids below were generated through mutation of an existing plasmid by PCR amplification of the template followed by Gibson assembly (one piece). See oligo list for sequences of PCR oligos.

| Plasmid | Template | PCR Oligos |
|---|---|---|
| pEP213 | pGB106 | oEP973/oEP974 |
| pEP218 | pGH504 | oEP977/oEP978 |
| pEP220 | pEP223 | oEP1031/oEP1034 |
| pEP221 | pEP223 | oEP1031/oEP1035 |
| pEP239 | pEP223 | oEP1051/oEP1031 |
| pEP241 | pEP223 | oEP1032/oEP1036 |
| pEP242 | pEP223 | oEP1010/oEP1011 |
| pEP243 | pEP223 | oEP1041/oEP1042 |
| pEP244 | pGB106 | oGB24/oGB33 |
| pEP245 | pEP223 | oEP1037/oEP1038 |
| pEP246 | pGB104 | oGB24/oGB33 |
| pGB86 | pGB76 | oGB175/oGB180 |
| pGB103 | pGB51 | oGB172/oGB173 |
| pGB104 | pGH419 | oGB34/oGB35 |
| pGB106 | pGB103 | oGB34/oGB35 |
| pGH419 | pGH424 | oGH367/oGH369 |

The plasmids below were generated through Gibson assembly reactions comprising multiple fragments. The strategy is detailed for each plasmid.

pEP1

The backbone of pET21b was amplified by PCR using oligos oEP318/oEP324. The resulting PCR product was combined with IDT gBlock gbEP11 in a Gibson assembly reaction, yielding pEP1.

pEP57

The MosSCI targeting plasmid backbone for *cxTi10816*, including the *C. elegans dpy-30* promoter, an N-terminal tagRFP-T and the *C. elegans unc-54* 3'UTR was amplified by PCR from pGH486 using oligos oGH1011/oGH1012. The DNA encoding amino acids 1-163 of *C. elegans* NECAP was amplified by PCR from cloned cDNA using oligos oEP366/oEP369. These two PCR products were combined in a Gibson assembly reaction, yielding pEP57.

pEP223

The pET-21b backbone along with a TEV cleavable HaloTag was amplified by PCR from pGB63 using oligos oEP1031/oEP1032. The DNA encoding amino acids 1-170 of human NECAP2 was amplified by PCR from pEP1 using oligos oEP1033/oEP641. These two PCR products were combined with IDT gBlock gbEP42 in a Gibson assembly reaction, yielding pEP223.

**Sequences of synthesized gene fragments**

gbEP11: Human NECAP2(1-170) for pET-21b

TTTAAGAAGGAGATATACATATGGAAGAAAGCGGCTATGAAAGCGTTCTGTG
TGTTAAACCGGATGTTCATGTTTATCGTATTCCGCCTCGTGCAACCAATCGTG
GTTATCGTGCAGCAGAATGGCAGCTGGATCAGCCGAGCTGGTCAGGTCGTCT
GCGTATTACCGCAAAAGGTCAGATGGCATATATCAAACTGGAAGATCGTACC
AGCGGTGAACTGTTTGCACAGGCACCGGTTGATCAGTTTCCGGGTACAGCAG
TTGAAAGCGTGACCGATAGCAGCCGTTATTTTGTTATTCGTATTGAAGATGGT
AATGGTCGCCGTGCCTTTATTGGTATTGGTTTTGGTGATCGTGGTGATGCCTTT
GATTTTAATGTTGCACTGCAGGATCACTTTAAATGGGTTAAACAGCAGTGCG
AATTTGCAAAACAGGCACAGAATCCGGATCAGGGTCCGAAACTGGATCTGGG
TTTTAAAGAAGGTCAGACCATCAAACTGAACATTGCCAACATGAAAAAAAAA
GAAGGCCATCATCACCATCACCACTAACCGCTGAGCAATAACTAGCATAACC
C

gbEP42: Human NECAP2(171-263) for pET-21b

AACATGAAAAAAAAAGAAGGCGCAGCGGGTAACCCCCGTGTACGTCCGGCG
TCTACTGGCGGCCTTTCCCTTTTGCCTCCACCACCGGGTGGAAAACCTCTAC
ACTTATTCCTCCACCAGGGGAACAACTGGCGGTAGGTGGCTCTTTGGTTCAGC
CTGCTGTGGCCCCTAGCAGTGGTGGTGCTCCAGTACCTTGGCCCCAACCGAA
CCCCGCTACCGCGGATATCTGGGGAGACTTCACCAAGTCCACGGGCTCTACC
TCAAGTCAGACGCAGCCAGGCACGGGCTGGGTACAATTTCATCACCATCATC
ACCACTGA

### Sequences of DNA oligos

| Oligo ID | SEQUENCE |
|---|---|
| oEP318: | CATCATCACCATCACCACTAA |
| oEP324: | ATGTATATCTCCTTCTTAAAGTTAAACAAAATTAT |
| oEP366: | AACGGGCGGTAGTGGAGGCACTGGTATGGGAGATTACGAGAACGTTTTAATG |
| oEP369: | TATCACCACTTTGTACAAGAAAGCTGGGTCTAACTTTTATCCTTTTTTCCAATGTTAATT |
| oEP512: | AGTACTAGCGGTGGCAGTGGAGGTACCGGCGGAAGCATGGTCAGCAAGGGAGAGGCAGTT |
| oEP513: | CCGTCTTTATAGTCACCATCGTGGTCTTTGTAGTCCTTGTAGAGCTCGTCCATTCCTCCG |
| oEP519: | TTCCGACTAAAAATCCCCAAATTTTCAGAGATTTCAGTACTAGCGGTGGCAGTGGAGGTA |
| oEP520: | AGTTGTACGGAGAAGAAAGATGACGTCATCGCTTATCCTCCTTTGTCGTCATCATCCTTA |
| oEP582: | GGAGGAACGGGCGGTAGTGGAGGCACTGGTTAGACCTAGCTTTCTTGTACAAAGTGGTGA |
| oEP584: | GGAGGAACGGGCGGTAGTGGAGGCACTGGTGCGGAACTGGAAAAACAGGATCTTTCTGCC |
| oEP641: | GCCTTCTTTTTTTTTCATGTTGG |
| oEP655: | AAAAAGTCGATAGAGAAGGCTTCAACACAC |
| oEP656: | TCGGTCAAATTTCCCGGTTTTTAAC |
| oEP657: | ACCCGATTTTCTCGGTTTTTCTCTC |
| oEP659: | ATAAAAGTACGGATTTTTGCTCGAAATCAAC |
| oEP660: | GGCCAATTTTGAGGATTCTTTGGC |
| oEP661: | TTTAGACTGAAAATTCCGATTTTTGAGCC |
| oEP662: | GGTGGAGAGAGAGAAGTGAAGAGACGC |
| oEP670: | GCAAACTGGGGCACAAACTTAATTCC |
| oEP793: | GATCACTTTCGTTATATCGAACGAAGCTAATTCTTGTACAAAGTGGTGATATCTGAGCTC |
| oEP808: | AACAACATGAAGTGGCAGTCGC |
| oEP812: | GGAGCACAGGGAGAAAGAGC |
| oEP894: | AGGCATTCGTGGGATGCGGGTTTCAAGAAGAGGGAGATGCTTTTGACTTTAATGTCACAC |
| oEP865: | GGATGCGGGTTTCAAGAAGAG |
| oEP969: | CGTCAAGTGAAGCTGATCCGTGGCGTAGTTGGCTCACTTGGATTAATCGTAATCCC |
| oEP970: | GGGATTACGATTAATCCAAGTGAGCCAACTACGCCACGGATCAGCTTCACTTGACG |
| oEP971: | CGTCAAGTGAAGCTGATCCGTGGCGTAGTTGGCTCACTTGGATTAATCGTAATCCCGCTA |
| oEP972 | TAGCGGGATTACGATTAATCCAAGTGAGCCAACTACGCCACGGATCAGCTTCACTTGACG |

| | |
|---|---|
| oEP973: | TGCTCGAGCACCTATGATTTGGATTGTAGGAGAATATGC |
| oEP974: | TAGGTGCTCGAGCATCGGGTTCATCCAG |
| oEP977: | ATTGTCGACAATCGCGATGATGTGCAG |
| oEP978: | CGATTGTCGACAATCTGGATGACGCG |
| oEP985: | TGCCGGTCCAAGCTTGGATTTAGCATTTGCAGCTGCACAAACCATCTCAATTAACATTGG |
| oEP987: | GTTATTATTTCGGATTGTCGACAATAGGGAGGACGTTCAAGGATACGCAGCAAAGACTGT |
| oEP1010: | ACCGATAACAGCCGTTATTTGTTATTCGTA |
| oEP1011: | ACGGCTGTTATCGGTCACGCT |
| oEP1014: | GGGACAAACCATCTCAATTAACATTGGAAAAAAAGACAAGTCATAGACTCAGCTTTCTTGTACAA AGTGGTGATATCTGA |
| oEP1016: | CAGTGAAAAGTTCTTCTCCTTTACT |
| oEP1020: | TAGTGCCACTGCTTCCACCGCCGCCAGGTGCCGGCTAAACCCAGCTTTCTTGTACAAAGT |
| oEP1031: | CATCACCATCATCACCACTGAGATC |
| oEP1032: | CATGCTTCCGCCGGTAC |
| oEP1033: | AGGTACCGGCGGAAGCATGGAAGAAAGCGGCTATG |
| oEP1034: | TCAGTGGTGATGATGGTGATGGCCTTCTTTTTTTTCATGTTGGC |
| oEP1035: | TCAGTGGTGATGATGGTGATGCTGCTGTTTAACCCATTTAAAGTG |
| oEP1036 | GGTACCGGCGGAAGCTGCGAATTTGCAAAACAGGC |
| oEP1037: | TGATGAGGGTGATGCCTTTGATTTTAATG |
| oEP1038: | CACCCTCATCACCAAAACCAATACCAAT |
| oEP1041: | TTTTGCAGCAGCTCAGACCATCAAACTGAACATTG |
| oEP1042: | CTGAGCTGCTGCAAAACCCAGATCCAGTTTCGG |
| oEP1051: | TCAGTGGTGATGATGGTGATGTTTCCCACCCGGTGGTG |
| oGB24: | CGCCGCCAGCCAATCTGCCCAGCCACCTGGCTGGTGATCTGGGACTGTTC |
| oGB26: | ATGAATAAGCCTCCGATCATCATATGTATATCTCCTTCTTATA |
| oGB27: | GCATTTATGAAACCCGCTGCTAATTAACCTAGGCTGCTGCCACCG |
| oGB28: | ATGATCGGAGGCTTATTCATCT |
| oGB29: | GCAGCGGGTTTCATAAATGCCA |
| oGB33: | GGGCAGATTGGCTGGCGGCGAGAAGGCATCAAGTA |
| oGB34: | AGCAAGAGTCTGGTGCCGCGCGGCAGCGGA |
| oGB35: | CTGCTTACCGCTGCCGCGCGGCACCAGACCTTGCTTGTTTCATCAGCTGTG |

oGB52: TGCATCACGGGAGATGCACT

oGB124: GGCTGGGTCCAGTTCCATCACCATCATCACCACTGA

oGB125: GAACTGGACCCAGCCGGTGC

oGB130: GGAGCAGTCACAAATCACGTCTCAAGTTGCCGGCCAAATTGGATGGCGTCGGGAGGGTAT

oGB147: ATCTCCCGTGATGCAGGGCCTGGCTCTTGGGGTAC

oGB148: TGTGAGTTTGCGAAACAAGC

oGB149: TTTCGCAAACTCACACATGCTTCCGCCGGTACCTC

oGB172: ATGATCGGAGGCTTATTCATCT

oGB173: TAGATGAATAAGCCTCCGATCATATGTATATCTCCTTCT

oGB175: CATCACCATCATCACCAC

oGB180: CAGTGGTGATGATGGTGATGGCTTCCGCCGGTACCTCCAC

oGH367: TAATGCTTAAGTCGAACAGAAAGTAATCG

oGH369: TCTGTTCGACTTAAGCATTATTTGCGATGAATCCCATGAC

oGH408: GCATTTTTCACATTTTCTAACATTTTTTCTGTTGAAAAG

oGH409: GCACATTTTAAGTCTGTAAAAGTGAAAACCCA

oGH411: TCCTATGCTCAGTCAGTGTATGAGC

oGH412: GCTTTTGGAGCATTTTGTTTTCTAATTTTGAATGA

oGH413: AAGTTTTATACCAAGTTTAGAACATGGATTCGG

oGH414: CGGTTCTGCATGCAGTTGTCTG

oGH415: CCAAAAAAATGTATCTGAATAAGTAAAGCAAAGTGATTC

oGH416: GTCTTAACCAAAGAGCAACAACAATACCT

oGH417: ACTATAACTTTTGATTGTTTTGTCAACAGCTAGC

oGH418: CCACTTTTTCTAATATTTCAAACTTGTGCTCGA

oGH419: TGTAAAAGTGGAGAGATGGCACGG

oGH430: GTTTTGCAAAGATATTTTAATGAAGTTTGGCTCA

oGH432: AAATTAATTGTTTCTACAGAGTGTTTCAATGTTTGAAC

oGH441: GCTCCAATTTCCTTGAAACCTCG

oGH442: CCTTGAAAGCTTTTTTTAAGTTTTTTAGGTG

oGH443: GATTTTTCAAAATTTTTAACATCGAAACTCCC

oGH444: GCCCGATTTTACAGGAACTCC

oGH445: CTAAAATTCTAAACTACAAAATAATAATAAAAATATC

oGH446: TGCAATTTTTACAGGTCAGG

oGH447: CTCGGAAATTCAAATTATACATCAAAAATTATCAC

oGH448: GAAATTCAGAATTATTTAGGGGAAAAGGC

oGH452: CCATTCATATTTTGTCTCAGGAGAATAC

oGH679: AGGTATTCAGACATTTTTCAAATGAAAATCTAC

oGH847: CCAAACTGAAGGTCAAGGTGGTC

oGH848: CCTTGACCTTCAGTTTGGTGCGC

oGH1014: CCGCCGTCGTTCTCTCCACCG

RWB099: GGCCTCCTTCGTCGTCTTCAGGATCCAATTCGAGCTCGAACAACAAC
